# Unexpected sounds non-selectively inhibit active visual stimulus representations

**DOI:** 10.1101/2020.04.16.044966

**Authors:** Cheol Soh, Jan R. Wessel

## Abstract

The brain’s capacity to process unexpected events is key to cognitive flexibility. The most well-known effect of unexpected events is the interruption of attentional engagement (distraction). We tested whether unexpected events interrupt attentional representations by activating a neural mechanism for inhibitory control. This mechanism is most well-characterized within the motor system. However, recent work showed that it is automatically activated by unexpected events and can explain some of their non-motor effects (e.g., on working memory representations). Here, human participants attended to lateralized flickering visual stimuli, producing steady-state visual evoked potentials (SSVEP) in the scalp-electroencephalogram. After unexpected sounds, the SSVEP was rapidly suppressed. Using a functional localizer (stop-signal) task and independent component analysis, we then identified a fronto-central EEG source whose activity indexes inhibitory motor control. Unexpected sounds in the SSVEP task also activated this source. Using single-trial analyses, we found that sub-components of this source differentially relate to sound-related SSVEP changes: while its N2 component predicted the subsequent suppression of the attended-stimulus SSVEP, the P3 component predicted the suppression of the SSVEP to the unattended stimulus. These results shed new light on the processes underlying fronto-central control signals and have implications for phenomena such as distraction and the attentional blink.

---

Unexpected perceptual events, such as sudden sounds, are known to disrupt ongoing thoughts and actions. For example, when a warehouse worker hears an unexpected glass-shattering sound while doing inventory, they may momentarily stop writing on a count sheet and indeed forget their count altogether. This type of stimulus-driven distraction produced by unexpected perceptual events is of key interest to researchers studying attentional capture and its control (Yantis, 1993; Simons, 2000; Corbetta et al., 2008; Awh et al., 2012). Cognitive psychology has generated substantial insights into how distractors affect attentional engagement with the same sensory domain (Bacon & Egeth, 1994; Theeuwes, 2004; Gaspelin et al., 2017; Liesefeld et al., 2017). Moreover, cognitive neuroscience has provided a comprehensive picture of neural activity after unexpected events (Courchesne et al., 1975; Corbetta & Shulman, 2002). However, relatively little is known about the exact neural mechanisms by which task-irrelevant unexpected events disrupt active, task-relevant attentional representations, especially across sensory domains. In a recent theoretical article, we proposed that unexpected sensory events engage an inhibitory brain mechanism that is capable of interrupting active neural representations that support active motor and non-motor (i.e., cognitive) functions (Wessel & Aron, 2017). At the core of this proposal is the assertion that a well-characterized neural mechanism for inhibitory control (which is well-known to be involved in suppressing active motor representations, and which is automatically triggered by unexpected events; Wessel & Aron, 2013; Wessel, 2018), can exert inhibitory influence even on non-motor representations, such as working memory or attention (Chiu & Egner, 2014, 2015; Wessel et al., 2016; Castiglione et al., 2019; Tempel et al., 2020).

The brain network in question implements its inhibitory function via a fronto-basal-ganglia (FBg) network that involves the pre-supplementary motor area (preSMA), the right inferior frontal cortex (rIFC), and the subthalamic nucleus (STN) of the basal ganglia (Nachev et al., 2007; Ridderinkhof et al., 2011; Schall & Godlove, 2012; Aron et al., 2014; Jahanshahi et al., 2015). Its inhibitory influence on movement is most well-studied in the stop-signal task (SST, Logan et al., 1984), where it implements the ‘stop’-component of the purported race between going and stopping (Aron et al., 2014). The temporal dynamics of this network can be non-invasively measured using scalp-electroencephalography (EEG), where its activity is indexed by a pronounced stop-signal induced fronto-central N2/P3 event-related potential (ERP) complex (de Jong et al., 1990; Kok et al., 2004; Huster et al., 2013; Wessel & Aron, 2015). The N2 component of this ERP complex has been proposed to reflect the detection of the stop-signal or the associated conflict (Schröger, 1993; Donkers & van Boxtel, 2004; Azizian et al., 2006; Enriquez-Geppert et al., 2010; Smith et al., 2010), whereas the P3 has been hypothesized to reflect the subsequently implemented inhibitory process (de Jong et al., 1990; Kok et al., 2004; Enriquez-Geppert et al., 2010; Wessel & Aron, 2015).

Our past work has shown that unexpected perceptual events automatically engage this same inhibitory mechanism (Wessel et al., 2012; Wessel & Aron, 2013, 2017), even when there is no explicit instruction to engage inhibitory control (Wessel, 2018a). Amongst other findings, this assertion is supported by the fact that unexpected events lead to a slowing of motor responses (Dawson et al., 1982; Parmentier et al., 2008) and elicit a fronto-central N2/P3 ERP complex that is morphologically similar to the fronto-central ERP complex elicited by stop-signals (Courchesne et al., 1975; Squires et al., 1975). Indeed, independent component analysis (ICA) of EEG data recorded in subjects that performed both the stop-signal task and tasks involving unexpected events suggests that both ERP complexes share a common underlying neural generator (Wessel & Aron, 2013; Wessel & Huber, 2019). Moreover, local field potential recordings from the human subthalamic nucleus (the key subcortical node of the inhibitory FBg-network) suggest that unexpected events engage this subcortical structure as well (Bočková et al., 2011; Wessel et al., 2016). In line with this, optogenetic inactivation of the STN negates the inhibitory influence that unexpected sounds have on behavior in mice. While unexpected sounds typically lead to a premature interruption of ongoing licking bouts, this effect is absent if the STN is inactivated, providing key causal evidence for the role of inhibitory control structures in surprise processing (Fife et al., 2017).

An important property of this inhibitory FBg mechanism is that it implements inhibition in non-selective, ‘global’ fashion, both after stop-signals and unexpected sensory events. This is most evident from experiments that use transcranial magnetic stimulation and electromyography to measure cortico-spinal excitability (for reviews, see Duque et al., 2017; Wessel & Aron, 2017; Derosiere et al., 2020). Such experiments show that the rapid reduction of cortico-spinal excitability that is found after stop-signals (Coxon et al., 2006; Stinear et al., 2009) extends even to task-unrelated motor effectors (Badry et al., 2009; Greenhouse et al., 2011; Cai et al., 2012; Goode et al., 2019). The same non-selective suppression of cortico-spinal excitability can be observed after unexpected events (Wessel & Aron, 2013; Dutra et al., 2018; see also: Novembre et al., 2018; Novembre et al., 2019). As mentioned above, we have proposed that this type of non-selective, ‘global’ suppression exerted by the inhibitory mechanism could explain why unexpected events have effects even on non-motor, cognitive representations (Wessel & Aron, 2017) – conceivably, if fronto-basal ganglia mediated inhibition is broad and non-selective enough, it may even affect non-motor representations (provided neural underpinnings of those representations are susceptible to this inhibitory circuit).

Indeed, some preliminary evidence for this proposal already exists. In a first series of studies, Chiu & Egner (2014) have found that pairing faces with the requirement to rapidly withhold a prepotent action – thereby triggering the inhibitory control network – inhibits the encoding of these face stimuli into memory. In a follow-up study, the strength of this effect related directly to the activation of the inhibitory FBg-network (Chiu & Egner, 2015). Further in line with this, both action-stopping in the stop-signal task and active suppression of memory contents (e.g., in the Think/NoThink paradigm, Anderson & Green, 2001) are accompanied by activity from the same neural source (Castiglione et al., 2019). Finally, in line with the finding that unexpected events engage the inhibitory network, we have found that the activity of the inhibitory FBg-network also mediates the disruptive effects of unexpected sounds on active verbal working memory representations (Wessel et al., 2016).

While these studies lend first preliminary support to the general idea that the FBg-network underlying motor inhibition could also explain the suppression of non-motor representations, all existing work so far is limited to mnemonic processes. Moreover, it has already been found that not all types of memory representations seem to be subject to the purported inhibitory influence of the FBg network (indeed, short-term visual memory representations as operationalized in the classic work of Vogel & Machizawa, 2004; Vogel et al., 2005, seem to be interrupted by other mechanisms, cf., Wessel, 2018a). Therefore, it is hitherto unclear which exact types of non-motor representations are potentially subject to interruption by inhibitory control exerted from the FBg-network, and whether its influence extends beyond the realm of mnemonic representations.

In the current study, we tested whether the activity of this mechanism could explain the effects of unexpected events on ongoing attentional representations. A highly influential body of past behavioral work indicates that indeed, attentional regulation may include inhibitory processes (Shapiro & Raymond, 1994; Klein & Taylor, 1994; Tipper et al., 1990). To test whether attentional representations are affected by inhibitory control signals after unexpected events, a novel task was designed in which participants attended to one of two concurrently presented rhythmic flickers. Such flickering visual stimuli are known to produce a steady-state visual evoked potential (SSVEP) – a stimulus-driven entrainment of parieto-occipital EEG activity to the frequency of the rhythmic sensory stimulation (Regan, 1989; Silberstein et al., 1995). Notably, the covert direction of attention towards a specific stimulus leads to an increase in the amplitude of the associated SSVEP (Morgan et al., 1996; Müller et al., 1998; Ding et al., 2006; Walter et al., 2012). In our task, we then presented unexpected sounds on a subset of trials while subjects were attending one of the flickering visual stimuli. We expected the unexpected sounds to rapidly and transiently reduce the amplitude of the SSVEP, reflecting an interruption of attentional engagement. In addition to this task, all participants also performed a stop-signal task, which served as a functional localizer for the FBg-network underlying inhibitory motor control. To test the hypothesis that this same neural mechanism is related to the interruption of attentional representations after unexpected events, we used ICA to identify the neural source signal underlying the N2/P3-complex in the stop-signal task and tested whether this source was also active following unexpected events (as found in prior work, cf. Wessel & Aron, 2013; Wessel & Huber, 2019). Finally, we then tested whether the activity of that EEG source related to the disruption of attention (i.e., the SSVEP) on a trial-to-trial basis.

## Materials and Methods

### Participants

In Experiment 1, 21 healthy adult college students (mean age: 19.05 years; SD: 1.12; three left-handed; 14 females) participated the experiment for course credit. In Experiment 2, 21 healthy adult college students (mean age: 20.52; SD: 2.14; one-left-handed; 11 females) participated. Six of those participants received course credit and the rest were compensated with $15 per hour. All participants had normal or corrected-to-normal vision. None of participants performed both experiments.

### Stimulus presentation

All stimuli were presented on a BenQ XL2420B 120Hz gaming monitor with 1ms response time, connected to an IBM compatible PC running Fedora Linux and MATLAB 2015b. Stimuli were presented using Psychtoolbox 3 (Brainard, 1997) at the monitor’s native resolution of 1920 x 1080 pixels. Responses were made using a standard QWERTY USB keyboard. Viewing distance was kept constant at 90 cm.

### Experimental paradigms

#### Experiment 1: Cross-modal SSVEP oddball task

The task (from here onwards referred to as the “SSVEP task”) was designed to induce sustained active perceptual / attentional visual representations during which unexpected sounds were presented on a subset of trials. All stimuli were presented in white color on a black background. A task diagram can be found in ***Figure 1***.

**Figure 1.**
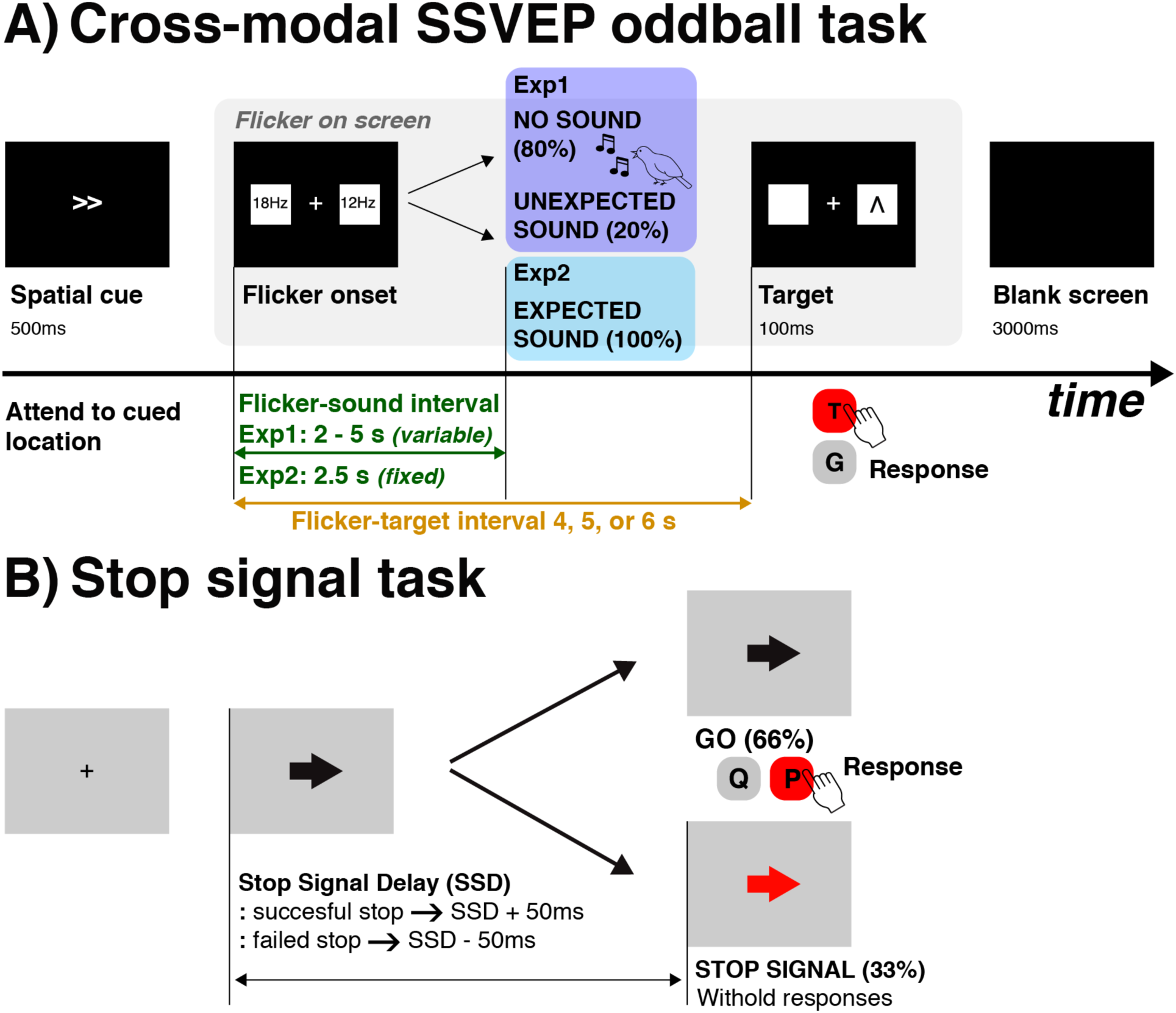
Task design. A) In Experiment 1, participants attended to a spatially cued rhythmic flicker (12 or 18 Hz) in order to detect a visual target that was superimposed on the cued flicker after a variable delay interval. On 20% of trials, unexpected sounds were presented in the delay interval, prior to target appearance. In Experiment 2, expected sounds were played 2.5 seconds after the flicker onset in all trials. B) In the stop signal task (performed by all subjects after the crossmodal SSVEP oddball task), participants made speeded responses to black arrows (go stimulus). On 33% of trials, the color of arrows changed into red (stop signal) after an adaptive delay, after which they were instructed to attempt to stop their response.

Each trial began with a centrally presented white double-arrow (<< or >>, 500ms duration, font size: 100) that informed the subjects which side of the display to covertly attend to. The initial double-arrow cue was followed by the SSVEP display, which consisted of a central fixation cross (+) flanked by two white flickering boxes (size: 9.66 × 9.66° of visual angle) that were presented to the left and right (offset 8.44° of visual angle laterally from center), with one box flickering at a frequency of 12Hz (f12) and the other flickering with a frequency of 18Hz (f18, half of the trials consisted of 12Hz left / 18Hz right displays, and the other half of 18Hz left / 12Hz right, presented in pseudorandom order). After a variable delay period, a visual target (∧ or ∨, font size 100) would appear in the center of the flickering box on the cued side of the display for 100ms. Participants were instructed to press either the ‘t’ (up, ∧) or ‘g’ (down, ∨) key on the keyboard to indicate the direction of the target stimulus. All responses were made using index or middle finger of the dominant hand (responses were made in vertical direction using the same hand to prevent potential cue-target spatial incompatibility effects that could result from the lateralized stimulus display). The flicker onset-to-visual-target delay was either 4, 5, or 6 seconds long, pseudo-randomly chosen from a uniform distribution of values. Participants were instructed to keep fixating on the central fixation cross while covertly attending to the cued flicker and monitoring it for the visual target. Horizontal eye movements were monitored by the experimenter between blocks (using the recording from the HEOG electrode on each trial) to ensure that the cued flickers were covertly attended and no overt saccades were made (trials with saccades to either side of the screen were removed during the analysis, see below). A blank screen was displayed for three seconds between trials.

On 20% of trials, unexpected sounds (unique bird song segments of 290 ms length) were played in the delay period between the flicker and the target onset (UNEXPECTED condition) through speakers positioned to either side of the computer screen. Subjects were not instructed to expect any sounds during the experiment, nor were sounds presented in the practice block. The inter-stimulus-interval between flicker onset and unexpected sound was drawn from a uniform distribution ranging from 2,000 to 5,000 msec, with the constraint that the chosen delay could not exceed the duration of the SSVEP display (i.e., the flicker onset-to-visual-target delay). In the remainder of trials (80%), no sounds were presented (NO SOUND condition). Sound volume was set at conversational level, which reliably evokes an orienting response without inducing a startle reflex. After one block of practice (24 blocks), participants performed 360 trials in total (36 trials per block, 10 blocks) with self-paced resting periods between blocks. Across the experiment, all conditions were counterbalanced (i.e., f12/f18 positions were equally distributed between left and right, as well as between UNEXPECTED and NO SOUND trials).

#### Experiment 2: Control SSVEP task

Experiment 1 was designed to test whether unexpected sounds disrupted the SSVEP by comparing the UNEXPECTED to the NO SOUND condition. We used Experiment 2 to confirm that the reduction of the SSVEP on UNEXPECTED trials in Experiment 1 was due to the unexpected nature of the sounds, and not due to the presence of a sound itself. The task was identical to Experiment 1, with the exception that *every* trial included a sound (a 600 Hz sine wave tone of 200ms duration), which was presented at the same time on each trial (exactly 2.5 seconds after flicker onset). Contrary to Experiment 1, participants were instructed to expect these sounds before the task, and their practice block (18 trials) included these sounds as well (hence, we will refer to this as the EXPECTED condition). To collect a matching number of trials in relation to the UNEXPECTED condition in Experiment 1 (and thereby equate the signal-to-noise ratio of the SSVEP), Experiment 2 contained only 72 trials (all within the EXPECTED condition), split evenly across two blocks.

#### Stop-signal task (functional localizer)

To evoke the neural signature of the inhibitory control mechanism, we used the same version of the SST that we used in prior work (Wessel, 2020). At the beginning of each trial, a black central fixation cross appeared for 500ms on a grey background. Then, a black arrow pointing to either the left or right was presented at the center of the display (go trials). Participants made speeded bimanual responses using either ‘q’ (left) or ‘p’ (right) key on the keyboard that spatially matched the go stimuli (e.g., ‘q’ for left arrow). In 33% of trials, the black arrows were replaced by red arrows (stop-signal) after certain amount of stop signal delay (SSD). The initial delay was set at 200ms. Participants were instructed to withhold responses on trials in which stop-signals appear. The SSD was adaptively adjusted in accordance with the stopping performance to ensure about 50% of probability in successful stopping [P(stop)], which is optimal for estimating the stop signal reaction time (SSRT) and guarantees motor prepotency on most successful stop-trials. The SSD was increased by 50ms after every successful stop; and decreased by 50ms after every failed stop. Experimenters instructed that making fast responses to the go stimuli and stopping responses to the stop-signals are both equally important. Verbal feedback was given after each block. Participants performed 300 trials (200 go, 100 stop), split evenly across five blocks.

#### Procedure

The SSVEP tasks in both experiments were always performed before the SST. The tasks were performed in this fixed order to avoid biasing participants towards using inhibitory control in the SSVEP task.

### Behavioral Analysis

All behavioral data from the SST were examined to check whether each subjects’ data conformed to the prediction of the race model of the stop-signal task (Logan et al., 1984). Specifically, we checked whether failed-stop trial reaction time was faster than Go-trial reaction time for each subject. We also checked whether the SSD algorithm converged around p(stop)=.5 by ensuring that the final p(stop) for each subject was between .4 and .6. SSRT was then computed using the revised version of the integration method with replacement of errors and misses, as suggested by Verbruggen et al. (2019).

### EEG Recording

We used a 62-channel electrode cap connected to Brain Vison MRplus amplifiers (BrainProducts) to record EEG at a sampling rate of 500 Hz. The reference electrode was Pz and ground was Fz. Two additional eye electrodes were placed beside and below the left eye to monitor for saccades and blinks, respectively.

### EEG preprocessing

EEG data were preprocessed using custom MATLAB scripts written in Version 2015b (TheMathWorks, Natick, MA). For each experiment, raw EEG data from the SSVEP task and the SST were imported into MATLAB and concatenated (i.e., the SST timeseries data were appended to the SSVEP task data). The merged timeseries were bandpass filtered (High-pass cutoff: .5 Hz; Low-pass cutoff: 50 Hz) using a Hamming-windowed sinc finite-impulse response filter (the default FIR filter in EEGLAB). All timeseries were visually inspected and non-stereotypical artifacts (muscle artifacts, transient electrode artifacts, etc.) were removed. Segments including saccades were manually removed and excluded from the further analyses to exclude trials in which attention was shifted overtly. Then data were re-referenced to the common average and entered into an infomax Independent Component Analysis (ICA) decomposition algorithm. Specifically, three different trial selections were performed prior to ICA, depending on which hypothesis was tested. Separate ICA solutions were generated for each of the three datasets.

1. To test the primary hypotheses (i.e., that unexpected sounds and stop-signals produce N2/P3 complexes from the same neural source, and that the activity of that source after unexpected sounds predicts the interruption of the SSVEP), all UNEXPECTED trials from the SSVEP task in Experiment 1 were combined with a matched amount of randomly selected NO SOUND trials, as well as the entire SST data. This was done to equate the signal-to-noise ratio of the SSVEP between UNEXPECTED and NO SOUND trials in the SSVEP task. Specifically, for each UNEXPECTED trial, a pseudo-event was generated within a randomly paired NO SOUND trial at the same after flicker onset at which the unexpected sound was played in the UNEXPECTED trial. Data from the SSVEP trials was included starting from 60ms prior to cue onset to 60ms following the response to the target.
2. To test whether EXPECTED sounds influence the SSVEP as well, data from the SSVEP portion of Experiment 2 were combined with the SST data for each of those subjects.
3. To test the attentional tuning of the SSVEP (i.e., to perform a manipulation check on the efficacy of the attentional cue), a dataset that only included the 288 NO SOUND trials from the SSVEP task (for the subjects in Experiment 1) or the 72 EXPECTED trials (for the subjects in Experiment 2) was generated.

Each of the resulting IC matrices for every subject was separately screened for stereotypic artifacts (e.g., blinks, EKG, channel noise), which were excluded prior to further analysis.

### Independent Component selection

#### Motor inhibition component selection

In line with previous work from us and others (Wessel, 2018b; Castiglione et al., 2019; Waller et al., 2019), one IC was selected from each participants’ ICA solution using the SST portion of the data as a functional localizer (this was only done for the ICA solutions generated to test the influence of the sounds on the SSVEP, and not on the ICA solution generated to test the SSVEP for attentional tuning effects, which did not include the SST data). In the following, this component will be referred to as motor inhibition independent component (MI-IC). The MI-IC shows four primary characteristics in the SST that have been demonstrated in our previous work (Wessel & Aron, 2015; Wessel et al., 2016; Wessel, 2017). First, the MI-IC shows maximal weights around fronto-central electrodes (FCz, Cz). Second, the MI-IC shows a pronounced positive deflection in its ERP, which peaks around 250-300 ms after stop-signals (the stop-signal P3), which is not present during matched time periods on Go-trials. Third, the onset of this ERP in the MI-IC occurs significantly earlier in successful stop-trials compared to failed stop-trials. This characteristic reflects a key prediction of the in the independent race model of the SST (Logan & Cowan, 1984), which holds that a faster stop process will lead to successful stopping. Fourth and finally, the onset of stop-related P3 is positively correlated to the behavioral measure of stopping speed (SSRT) across subjects, such that subjects with an earlier onset of the P3 component in the MI-IC have a shorter SSRT (for details, cf. Wessel & Aron, 2015).

To extract the IC for each subject that most closely corresponded to these criteria, we first selected those ICs that showed scalp topographies with maximal weights at fronto-central electrodes (F1, Fz, F2, FC1, FCz, FC2, C1, Cz, C2). Second, the resulting ICs were individually backprojected into channel space and their fronto-central stop-trial ERP was plotted to ensure that they showed a fronto-central N2/P3 complex following stop-signals. The relationship between the activity of these components and stopping behavior was then validated as follows.

#### Motor inhibition component validation

To identify the onset of the stop-signal P3 feature of the MI-IC, four types of trials in the SST portion of each subjects’ data were investigated: successful stop (SS) and matched go (SGo); failed stop (FS) and matched go trials (FGo). Go-trials were matched to stop-trials by selecting one go-trial per stop-trial in which the SSD staircase was at the same point (i.e., for a stop trial with an SSD of 200ms, we selected a go trial on which a stop-signal would have appeared at 200ms, had there been one). We then compared the mean sample-to-sample difference in MI-IC activity between stop and matched go-trials (SS vs. SGo; FS vs. FGo) within each subject using label-switching permutation testing (10,000 iterations, p = .01, corrected for multiple comparisons using the false-discovery rate method, FDR, Benjamini et al., 2006). The onset of the P3 was then defined as the first sample at which stop and matched go-trial MI-IC ERPs significantly diverged prior to the peak of the P3 (in essence, the peak of the P3 was identified, and the algorithm then worked ‘backwards’ towards the stop-signal until the stop-vs-go difference was no longer significant). The thusly identified P3 onset was then compared between successful and failed stop-trials across subjects using a paired-samples t-test. Moreover, the onset of the P3 on successful stop-trials was correlated to each subjects’ SSRT estimate using Pearson’s correlation coefficient. These procedures are identical to our first report of these properties (Wessel & Aron, 2015).

#### SSVEP component selection

Independent components reflecting the SSVEP were identified based on topographical and frequency criteria for all three ICA solutions for each subject. To be selected as an SSVEP component, an IC had to fulfill the following criteria: First, it had to show weight matrix maximum at parieto-occipital electrodes (PO8, PO7, PO4, PO3, P8, P7, P6, P5, P4, P3, P2, P1, O2, and O1). Second, it had to be among the top eight ICs in terms of explained variance of the whole-scalp 12 and 18 Hz response (identified by EEGLAB’s built-in spectopo() function). This resulted in an average of 3.24 components per subject that were selected as SSVEP components (range: 2-6).

#### Manipulation check: Unexpected events and stop-signals elicit N2/P3 complexes in the same IC

After selecting the MI-IC and confirming its properties in the SST, we then aimed to replicate prior findings showing that unexpected events evoke an N2/P3 complex within that same neural source (Wessel & Aron, 2013; Wessel & Huber, 2019). To this end, the MI-IC was back-projected into channel-space, and the fronto-central ERP (average at FCz and Cz) of that back-projection was time-locked to the onsets of UNEXPECTED sounds in the SSVEP task and the above-mentioned ‘pseudo-events’ on NO SOUND trials in Experiment 1 (−500 to 1000ms), as well as to the EXPECTED sounds in Experiment 2. We then compared the subject-average activity time-course using sample-to-sample t-tests in the post-event period. Specifically, a paired-samples test was used to test the difference between UNEXPECTED sounds and the NO SOUND condition in Experiment 1, and an independent samples t-test was used to test the difference between the UNEXPECTED sounds in Experiment 1 and the EXPECTED sounds in Experiment 2. Both resulting vectors of p-values were corrected for multiple comparisons using the FDR-method to a critical alpha-level of .05.

### Time-frequency analysis

To convert the time-domain EEG signal to the time-frequency domain, the entire EEG timeseries were bandpass filtered with 30 linearly spaced center frequencies spanning 1 – 30 Hz with a range of 1 Hz around the respective center frequencies. The analytical amplitude of the signal at each center frequency was then computed using the square of the absolute of the Hilbert coefficients, identified using MATLAB’s hilbert() function.

### Steady-state visual evoked potential analysis

#### SSVEP extraction

All SSVEP activity was quantified from the time-frequency time series at electrodes PO7 (left hemisphere) and PO8 (right hemisphere). For frequency and attentional tuning analyses, EEG data with all ICs were used to match prior studies. The remainder of SSVEP analyses used the backprojection of the EEG data produced using the SSVEP ICs because our main hypothesis tested how neural activities from two statistically independent neural sources (MI-IC and SSVEP IC) interact with each other after unexpected sounds. The IC-based source-signal approach not only avoids cross-contamination of channel-space activity due to volume conduction, but it also increases the single-trial signal-to-noise ratio of both the SSVEP and the MI-IC activity. Five participants from Experiment 2 were excluded from the SSVEP analyses because after artifact rejection, at least one of their SSVEP conditions included fewer than 10 trials (Experiment 2 contained only 18 trials per each of the four SSVEP conditions).

#### Manipulation check: frequency tuning

To identify whether there was an SSVEP entrained to the visual stimuli, the data were segmented from -300 to 3,000 ms relative to flicker onset. Each trial was then baseline corrected by converting the amplitude to a z-score relative to the 300ms pre-stimulus period. For each trial, we then computed the median amplitude of the z-scored time-frequency amplitude at both 12 and 18Hz from the contralateral hemisphere to the location of the f12 or f18 flicker. These values were then averaged to produce the trial-average SSVEP amplitude for each frequency (12/18Hz ERSP) contralateral to each flicker type. We then tested whether the SSVEP at either hemisphere was entrained more strongly to the frequency of the flicker in the contralateral visual field using paired-samples t-tests.

#### Manipulation check: Attentional tuning

We then investigated whether instructed shifts in covert attention increased the amplitude of the SSVEP, in line with previous literature (e.g., Regan, 1989; Müller et al., 1998; Ding et al., 2006). To this end, four SSVEP time series were investigated: 12 Hz attended, 12 Hz unattended, 18Hz attended, 18Hz unattended. These analyses were performed on the contralateral electrode only. To investigate the effect of attentional tuning over time, the z-scored single-trial data described above was binned into consecutive segments of 200ms, and the attended condition was tested against the unattended condition for each frequency using paired-samples t-tests.

#### Hypothesis test: SSVEP change after unexpected sounds

For all Experiment 1 datasets, each UNEXPECTED trial was paired with a matching NO SOUND trial as described above. Trials were then epoched into -500 to 1000ms segments around the sound for the UNEXPECTED trials, and around the same time point for the matching NO SOUND trial. For all Experiment 2 datasets, the data were time-locked to the EXPECTED sound. For all three trials types, both the attended (contralateral to cued location) and the unattended (ipsilateral) SSVEP were averaged across trials, and the resulting data were z-scored relative to the 500ms period prior to sound onset (or the ‘pseudo’-sound in case of the matched NO SOUND trials). Differences between the resulting average time-courses were then tested for significance using a sample-to-sample 2×2 ANOVA. Specifically, to test whether UNEXPECTED sounds reduced the SSVEP compared to the NO SOUND condition, we analyzed the data from Experiment 1 using the repeated-measures factors SOUND (unexpected vs. no sound) and ATTENTION (attended vs. unattended). Furthermore, to test whether any change after the UNEXPECTED sound was due to the expectancy violation, rather than presence of the sound itself, we compared the UNEXPECTED sound condition from Experiment 1 with the EXPECTED sound condition from Experiment 2 using the between-subject factor SOUND (Exp1: unexpected vs. Exp2: expected sound) crossed with the within-subject factor ATTENTION (attended vs. unattended). Both ANOVAs were applied to each sample point individually, resulting in three vectors of p-values (main effect of SOUND, main effect of ATTENTION, SOUND * ATTENTION interaction) for each analysis. These p-values were then corrected for multiple comparisons using the FDR-method to a critical alpha level of .05.

### Single trial general linear model: N2/P3 to SSVEP relationship

We tested our main hypothesis that interruptions of the SSVEP by unexpected sounds will be related to MI-IC activity triggered by those sounds using a single-trial GLM. To do so, we quantified the peak amplitude of the N2 and P3 portions of the N2/P3 complex in the MI-IC on in each trial with an UNEXPECTED sound in Experiment 1, as well as the reduction of the SSVEP on the same trial. The N2 peak amplitude was quantified by measuring the activity-minimum in the MI-IC backprojection at fronto-central electrodes FCz and Cz between 140 and 300ms following the time-locking event. The P3 peak amplitude was quantified by measuring the activity-maximum within a 150ms window starting from the peak latency of the N2 within that same time course.

To identify the change in trial-to-trial SSVEP after an unexpected sound on the same trials, we first conducted a group-level ANOVA on the trial averages. Specifically, in order to find a common time window for this analysis regardless of the type of SSVEP frequency, and to provide an independent contrast to identify this window, we conducted repeated-measures 2-way ANOVA with factor SOUND (unexpected vs. no sound) and FREQUENCY (12 Hz vs. 18Hz). As above, this was repeated for all samples and then FDR-corrected to reach a critical alpha level of .05. The time window in which the main effect of SOUND was significant was used to quantify the degree of SSVEP disruption at the single trial level. The SSVEP reduction was quantified as the change from baseline within that time window (38-744 ms).

Based on these values for the SSVEP ICs and the MI-ICs, four single-trial GLMs were generated for each participant, relating the single-trial amplitude of the MI-IC (N2 and P3) to the single-trial amplitude in the SSVEP (attended and unattended). Both predictors and DVs were standardized prior to the calculation of the coefficients. The resulting regression beta coefficients were Fisher’s z-transformed to ensure a normal distribution prior to statistical testing. The thusly-transformed beta-weights for each subject were then tested for significant differences from 0 using paired-samples t-tests.

### Temporal order of N2, P3, and SSVEP reduction

Finally, we compared the latencies of the ERP peaks in the MI-IC and the SSVEP reduction in the SSVEP-IC. These latencies were quantified on each UNEXPECTED trial and collapsed across all conditions to be averaged, and then compared across subjects using paired-samples t-tests. We predicted that the timing of N2 and P3 would reliably precede the SSVEP suppression.

## Results

### Stop-signal task (functional localizer)

#### Behavior

Consistent with the assumption of the independent race model (Logan & Cowan, 1984), all participants across both experiments showed slower go-trial RT (mean: 557.16 ms; SEM: 13.37) compared to failed stop-trial RT (mean: 482.53 ms; SEM: 11.30). The SSD staircase algorithm successfully kept the probability of stopping around .5, with a range from 0.47 to 0.57. Average SSRT was 216 ms (SEM: 5.14) and average SSD was 336 ms (SEM: 16.77). Average error rate was 0.38% and miss rate was 3.37%. Overall, these results represent a typical parameter range for healthy young adults.

#### MI-IC validation

Validating the MI-IC included testing whether 1) the MI-IC P3 onset on successful stop trials occurred reliably earlier than on failed stops; and 2) the P3 onset in successful stops was correlated to the behavioral measure of stopping speed (SSRT) across subjects. The N2/P3 complex in the MI-IC in the SST is depicted in ***Figure 2A. Figure 2B*** shows that P3 onset on successful stop-trials (mean: 258.43 ms; SEM: 6.51) was significantly earlier compared to failed stop-trials (mean: 284.71 ms; SEM: 7.59), t_(41)_=-6.26, p<.001. Finally, ***Figure 2C*** shows that the onset of the successful stop-trial P3 was positively correlated with SSRT (r = 0.52, p<.001). These results suggest that the P3 ERP component of the MI-IC index the activity of the inhibitory control process in the SST, as described in prior work.

**Figure 2.**
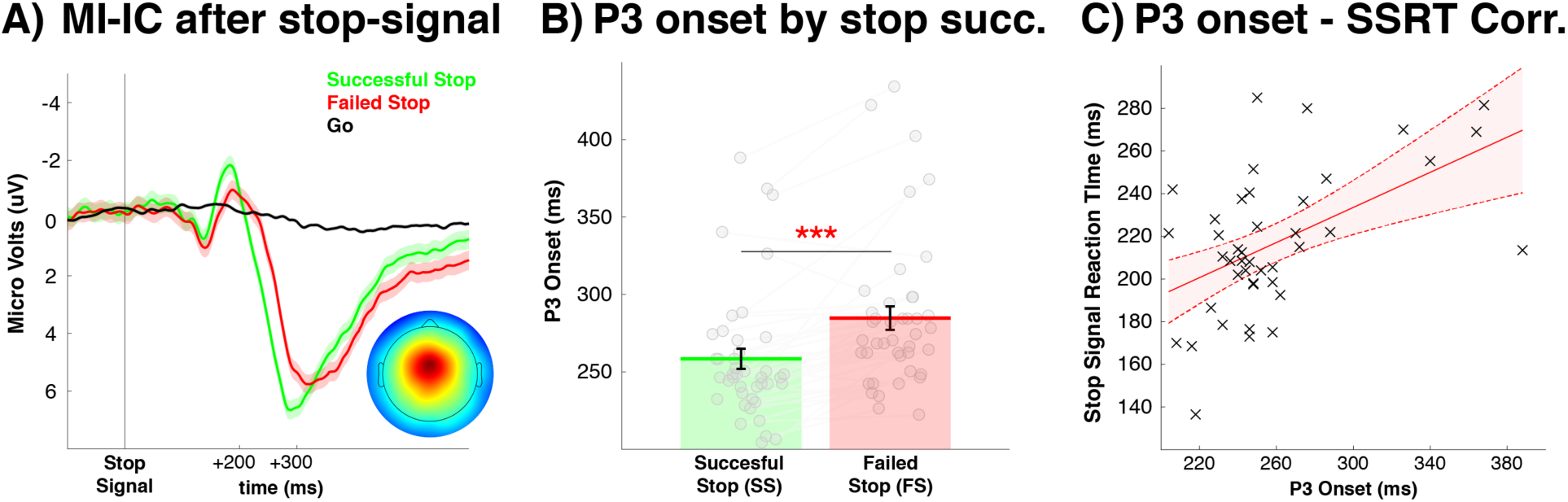
MI-IC activity after stop-signals. A) MI-IC fronto-central ERP time-locked to stop-signals in the SST data. Scalp topography inset represents averaged MI-IC weights from all participants. B) P3 onsets in successful (SS) vs. failed stop (FS) trials. C) Cross-subject correlation between SS P3 onset and SSRT.

#### Manipulation check: MI-IC activity following unexpected sounds

In line with our hypotheses and prior work, the neural source underlying the MI-IC also showed a pronounced N2/P3 following unexpected sounds in the SSVEP task (***Figure 3***).

**Figure 3.**
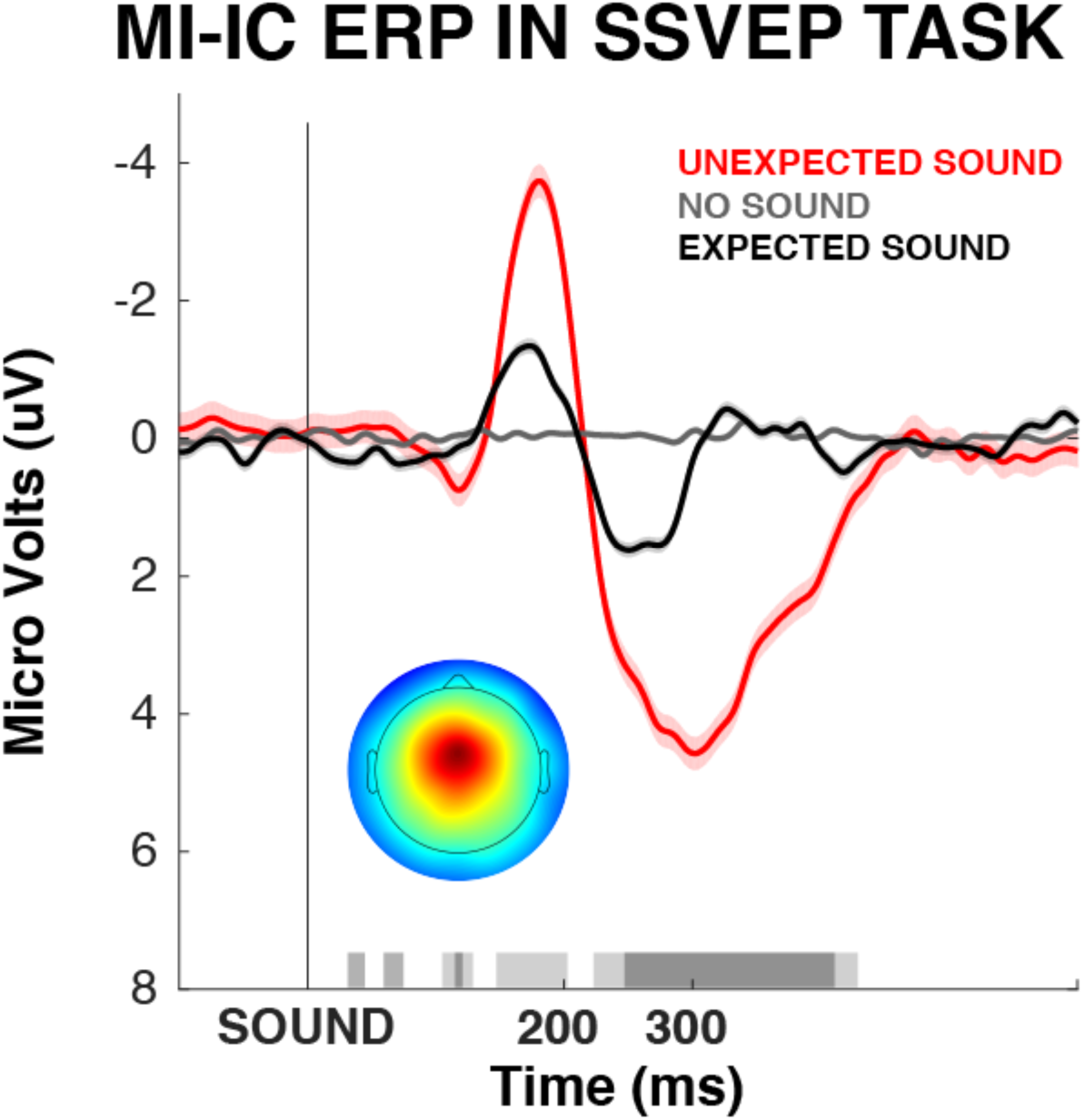
MI-IC fronto-central ERP activity after UNEXPECTED/EXPECTED/NO sound stimuli in the crossmodal SSVEP oddball task. Bright grey shade at the bottom of the figure indicates significant difference between UNEXPECTED vs. NO SOUND trials. Dark grey shade indicates significant difference between UNEXPECTED vs. EXPECTED sound trials.

### Steady-state visual evoked potentials

#### Manipulation check: frequency tuning

The hemisphere contralateral to f_12_ showed reliably higher ERSP_12Hz_ compared to ERSP_18Hz_ (t_(36)_=3.29, p<.01, Fig.4A), indicating that the 12Hz flickers successfully entrained a contralateral SSVEP. In turn, the hemisphere contralateral to f18 showed significantly higher ERSP_18Hz_ compared to ERSP_12Hz_ (t_(36)_=3.81, p<.001, Fig.4A), indicating that the 18Hz flickers successfully entrained a contralateral SSVEP as well.

**Figure 4.**
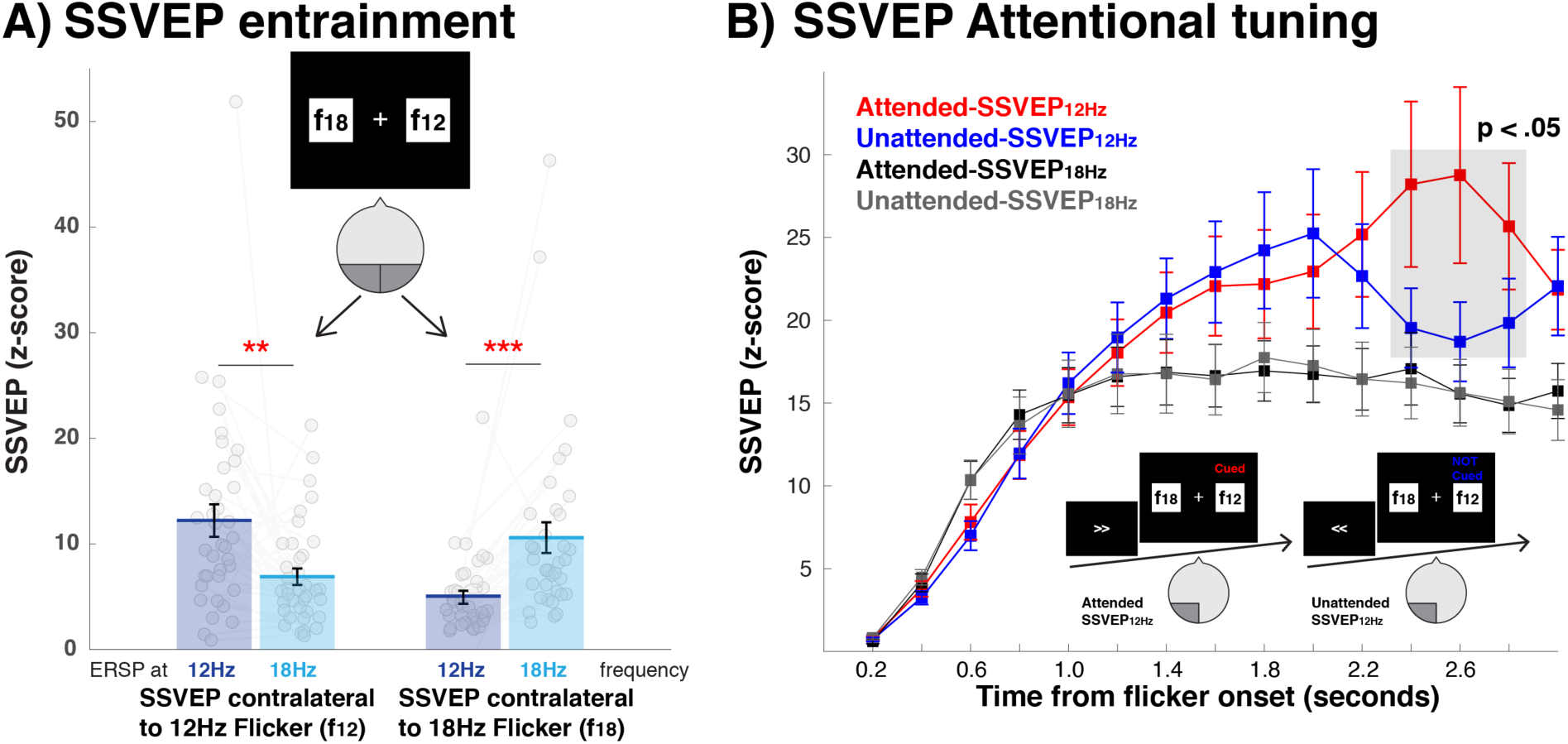
Frequency entrainment and attentional tuning effects in the SSVEP. A) Cross-frequency comparison of the SSVEP within the contralateral hemisphere. B) Binned time series after the flicker onset. Grey shade indicates significant difference between attended vs. ignored in the SSVEP to 12Hz flickers.

#### Manipulation check: Attentional tuning

As can be seen in ***Figure 4B***, we found a significant attentional enhancement effect in the time window from 2.2 to 2.8 seconds after flicker onsets (t_(36)_>2.1, p<.05), but only in the 12Hz condition. No such effect was found for the 18Hz condition. While this was not the expected outcome of this manipulation, this is in line with prior reports showing that attentional tuning of the SSVEP is often limited to the alpha range (Ding et al., 2006).

### Main hypotheses

#### SSVEP disruption by unexpected sounds

We then tested if unexpected sounds interrupted the ongoing SSVEP. A repeated-measures 2-way ANOVA with factor SOUND (unexpected vs. no sound) and ATTENTION (attended vs. unattended) showed main effects of SOUND from 0 to 770 ms in the 12Hz SSVEP (***Figure 5A***). Despite a visible reduction of the unexpected-sound SSVEP in the 18Hz SSVEP, there was no main effect of SOUND in the 18Hz SSVEP that survived corrections for multiple comparisons (***Figure 5B***). A mixed-model 2-way ANOVA with factor SOUND (Exp1: unexpected vs. Exp2: expected sound) and ATTENTION showed main effects of SOUND from 0 to 752 ms in the 12Hz SSVEP (***Figure 5C***); and from 148 to 616 ms in the 18Hz SSVEP (***Figure 5D***). There was no significant main effect of ATTENTION or SOUND*ATTENTION interaction for either ANOVA. This indicates that after unexpected sounds, the SSVEPs to both attended and unattended stimuli were significantly disrupted compared to no sound and expected sound trials.

**Figure 5.**
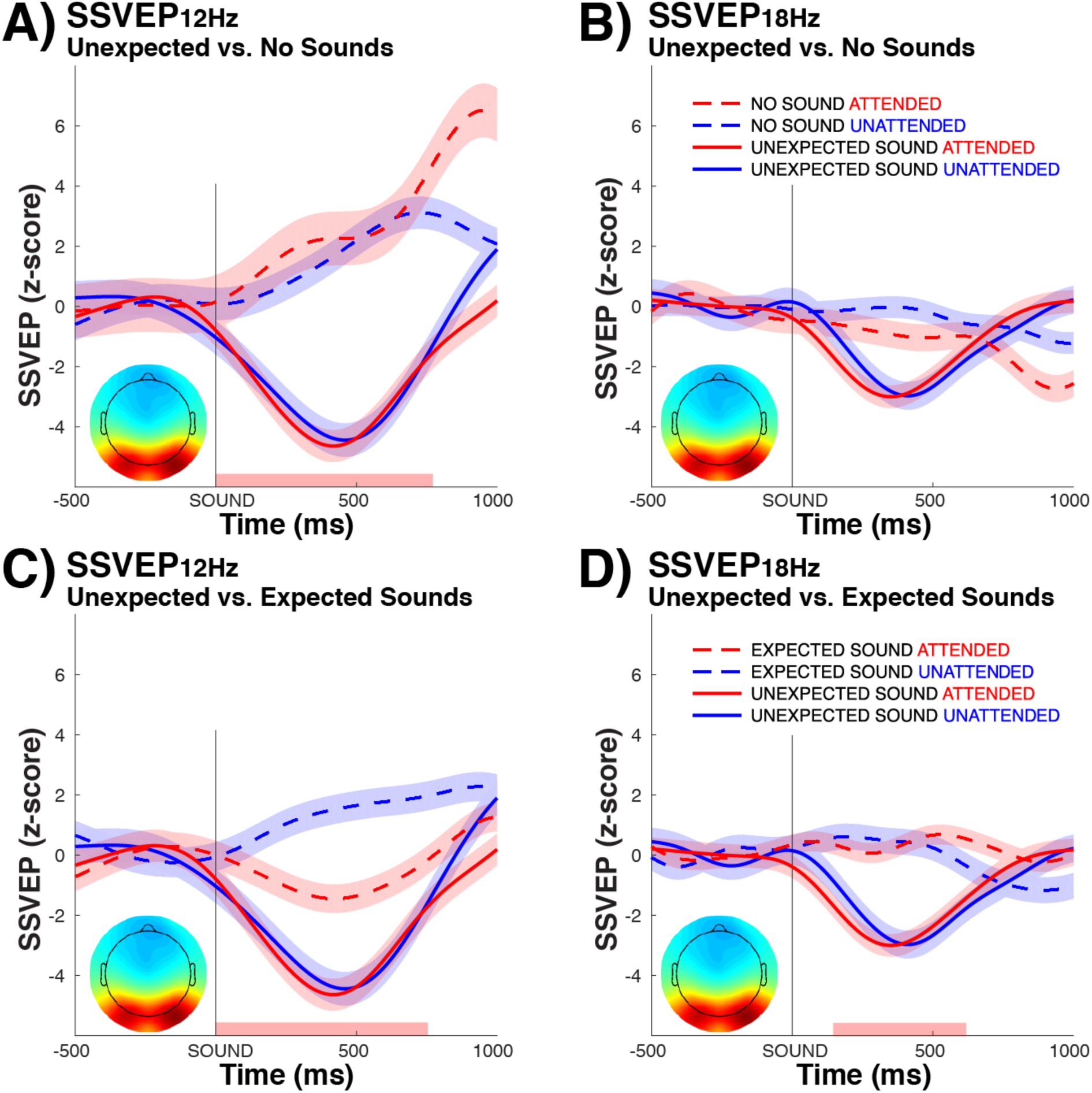
Suppressive effects of unexpected sounds on the active SSVEP. Time course of UNEXPECTED vs. NO SOUND trials in A) 12 Hz SSVEP; and B) 18Hz SSVEP. Time course of UNEXPECTED (Experiment 1) vs. EXPECTED sound (Experiment 2) trials in C) 12Hz SSVEP; and D) 18Hz SSVEP.

#### N2/P3 to SSVEP relationship

Our main hypothesis investigated the relationship between the activity of the MI-IC following unexpected sounds and the modulation of the SSVEP by those same sounds on the same trial. Since both N2 amplitudes and SSVEP reductions are negative-signed variables (-), greater N2 amplitudes leading to greater SSVEP decrements would result in positive beta weights, whereas the opposite would be true for the positive-valued P3 component (see ***Figure 6B*** for direction of each activity). We found that the MI-IC N2 and P3 were differentially related to the two components (attended and unattended) of the SSVEP IC. Specifically, the N2 amplitude reliably predicted the degree of suppression in the attended SSVEP (t_(20)_=2.67, p=.014, ***Figure 6A***), where the P3 amplitude reliably predicted the surprise-related decrement in the unattended SSVEP (t_(20)_=-2.44, p=.023, ***Figure 6A***).

**Figure 6.**
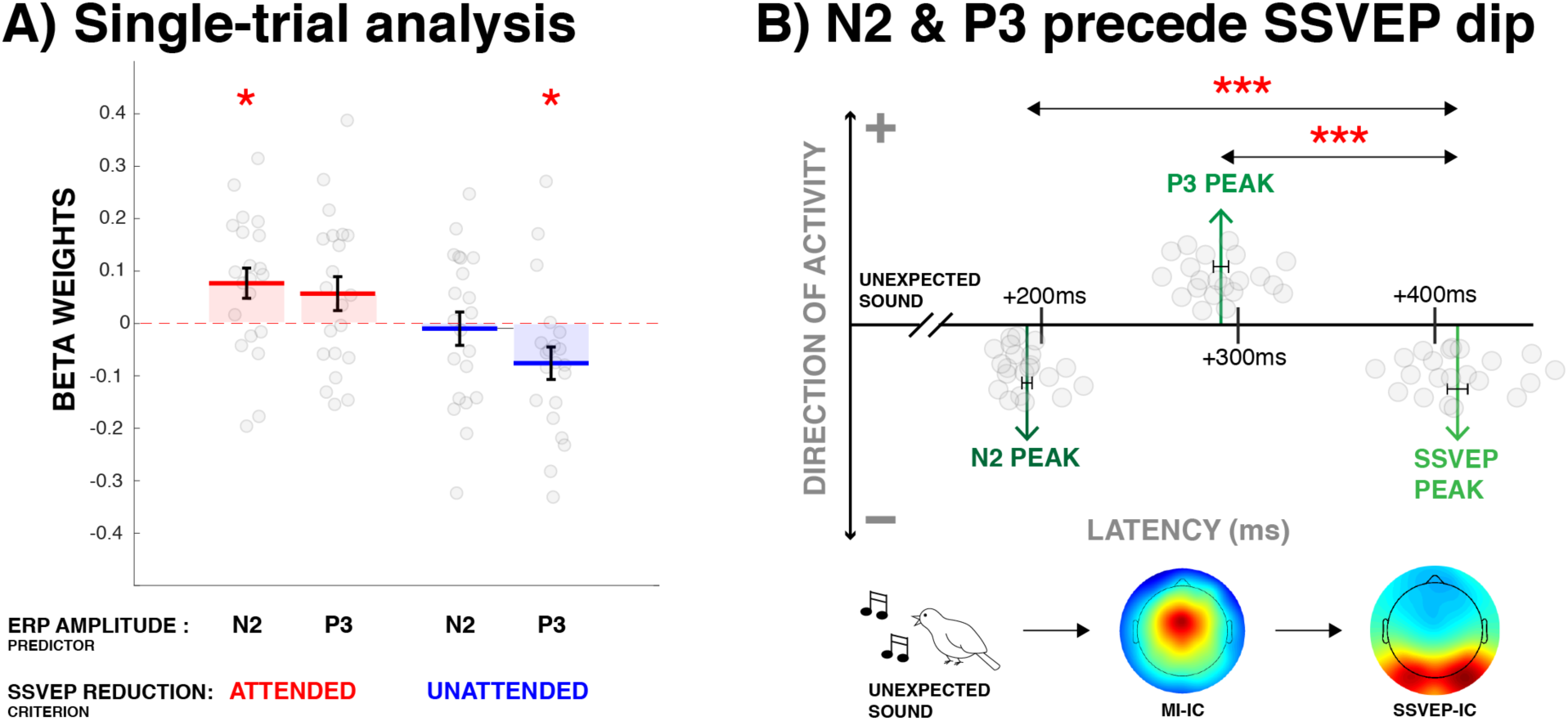
Single trial GLM results and peak onset comparison between MI-IC and SSVEP IC. A) Trial-to-trial relationship between N2/P3 amplitudes and SSVEP reduction to attended and unattended stimuli after unexpected sounds. B) The N2 and P3 peak latencies in the MI-IC backprojection and SSVEP suppression latencies in the SSVEP IC following unexpected sounds.

#### N2, P3, SSVEP reduction latencies

The average N2 peak latency was 194.98 ms (SEM: 2.56), whereas the average P3 peak latency was 292.71 ms (SEM: 3.82). The average latency of the SSVEP interruption was at 411.55 ms (SEM: 5.16). Both N2 (t_(20)_=-34.46, p<.0001) and P3 (t_(20)_=-17.73, p<.0001) latency were significantly earlier than the SSVEP latency (***Figure 6B***). These findings demonstrate that MI-IC activity following the unexpected sound preceded the suppression of the SSVEP.

## Discussion

In the current study, we investigated whether the interruption of visual attention after unexpected events is related to the activity of a well-known brain mechanism for inhibitory control. In a newly-developed paradigm, we first found that unexpected sounds lead to a suppression of SSVEP amplitudes to both attended and unattended visual stimuli. Moreover, a control experiment confirmed that this was not true following expected sounds. Using a functional localizer task to elicit the neural signature of a well-characterized brain mechanism for motor inhibition, we then replicated the finding that this EEG source (the MI-IC) was indeed active following unexpected sounds. Then, using a single-trial analysis of the independent components underlying inhibitory control and the SSVEP, we found that specific parts of the MI-IC response to unexpected sounds related to specific changes in the SSVEP on the same trial. Namely, the amplitude of the N2 potential of the fronto-central N2/P3-complex related to the suppression of the SSVEP to the attended stimulus, whereas the P3 potential related to the suppression of the SSVEP to the unattended stimulus.

These results provide new empirical evidence for the proposal that the brain’s inhibitory control mechanism is even broader than previously thought. Indeed, instead of solely affecting motor representations (e.g., during action-stopping in the stop-signal task), attentional interruptions after unexpected events appear to potentially result from inhibitory control as well. Indeed, the proposal that inhibitory control could affect non-motor representations goes back to the original work on the stop-signal task and the underlying race model, in which it was already proposed the stop-signal task invokes a mechanism that serves to “inhibit thought and action” (Logan et al., 1984). Notably, though, the vast majority of the subsequent work on this paradigm has focused on the stopping of action. In cognitive neuroscience, this work on action-stopping has firmly established a neural mechanism for motor inhibition, which serves to suppress ongoing motor representations when necessary (for review, cf. Verbruggen & Logan, 2009; Levy & Wagner, 2011; Ridderinkhof et al., 2011; Aron et al., 2014; Verbruggen et al., 2019). Only recently have cognitive neuroscience studies begun to harken back to the “thought” part of Logan and colleagues original proposal, thereby extending the effective range of this inhibitory mechanism to non-motor representations. However, these studies have so far exclusively focused on mnemonic representations, including short-term memory for face stimuli (Chiu & Egner, 2014, 2015), verbal working memory (Wessel et al., 2016; Castiglione et al., 2019), and motor sequence memory (Tempel et al., 2020). Expanding on this work, our study is the first of its kind to demonstrate that active attentional representations could be subject to the same type of inhibition as mnemonic and motor representations, mediated via the same neural pathway.

In our previous theoretical work on this topic (cf., Wessel & Aron, 2017), we have argued that the neuroanatomy of the neural pathway underlying inhibitory control could offer an explanation as to why non-motor representations like memory and attention could be subject to the same type of inhibition as the motor representations. The mechanism underlying inhibitory control involves a well-specific network of cortical and basal ganglia regions (Aron et al., 2007; Wiecki & Frank, 2013; Jahanshahi et al., 2015; Chen et al., 2020). Mechanistically, it is thought that the cortical areas of this network (which include the areas that produce the N2/P3 complex; Enriquez-Geppert et al., 2010; Huster et al., 2012) signal the need to initiate inhibitory control to the basal ganglia; specifically, to the subthalamic nucleus (Swann et al., 2011; Ray et al., 2012; Schmidt et al., 2013). In turn, the subthalamic nucleus is then thought to interrupt the thalamo-cortical loops that are underlying active motor representations (via the output nuclei of the basal ganglia, most notably the globus pallidus, Alexander & Crutcher, 1990; Nambu, 2008; Tanibuchi et al., 2009; Goldberg et al., 2013). Within that same framework, we propose that such fronto-subthalamic-pallidal-thalamocortical inhibition could potentially extend to *any* type of active neural representation that is maintained via thalamocortical loops (Wessel & Aron, 2017). Indeed, of core relevance to the current finding is the fact that the nuclei of the thalamus are a key nodes in the maintenance of not just motor representations, but also of active attentional representations (e.g., Desimone et al., 1990; McAlonan et al., 2008). In fact, while classic conceptualizations thought of the thalamus as merely a relay of sensory information, more recent work has found that thalamic activity exerts gain control over attentional representations, especially in the visual system (e.g., Saalmann & Kastner, 2009; Wimmer et al., 2015; Mease et al., 2016) and that lesions to the thalamus crucially interfere with attentional selection (Snow et al., 2009). If sustained visual attention, such as the type that is operationalized in our current paradigm, is indeed dependent on thalamocortical loops, it is conceivable that the same type of inhibitory influence from the basal ganglia that regulates motor behavior could also function to rapidly inhibit these active attentional representations.

In addition to this hypothesized subcortical overlap between the neural networks that regulate motoric and attentional representations, it is notable that the *cortical* areas of the fronto-basal ganglia inhibitory control network also overlap substantially with the wider networks implicated in attentional control in general. Indeed, Corbetta & Shulman’s seminal account of the ventral attention network – which ostensibly functions as a ‘circuit breaker’ that is triggered by suddenly appearing, behaviorally relevant stimuli (Corbetta & Shulman, 2002; Corbetta et al., 2008) – includes both cortical areas of the proposed fronto-basal ganglia inhibitory control network (the preSMA and the rIFG). Notably, however, in its original conceptualization, this purported ventral attention network does not include any specific areas in the basal ganglia, which we would propose based on our circuit model of inhibitory control. However, the absence of prominent basal ganglia involvement in the work on the ventral attention network may be a consequence of the fact that most of the work on that network has been performed using functional magnetic resonance imaging at field strengths that lack a sufficient amount of signal to noise ratio in small subcortical structures (Forstmann et al., 2017), especially in the subthalamic nucleus (de Hollander et al., 2017). Therefore, while it is still unclear how attention may be regulated using subcortical circuitry outside of the thalamus, it is possible that the type of attentional orienting implemented by the ventral attention network is indeed aided by an active inhibitory effort that suppresses ongoing attentional representations, implemented by the specific regions that form the inhibitory FBg-network (Wessel & Aron, 2017). Hence, future studies could use the current paradigm to study the involvement of the basal ganglia in the interruption of active attentional representations by unexpected sensory events.

In this vein, it is important to mention that the scalp-EEG methods used here do not allow any inferences about such underlying specific cortical or subcortical circuitry (though notably, the trial-to-trial variance of N2/P3 complex is correlated with BOLD activity in cortical areas that belong to both the fronto-basal ganglia inhibitory control network and the ventral attention network, Enriquez-Geppert et al., 2010; Huster et al., 2012). However, scalp-EEG does provide a temporally precise picture of the activity of the overall network. In this respect, the fronto-central N2/P3 complex is well-studied during both action-stopping and surprise processing (for reviews, see Polich, 2007; Folstein & Van Petten, 2008; Huster et al., 2013; Kenemans, 2015). However, in both literatures, the respective interpretation of the two constituent components of this complex waveform (the N2 and the P3) is still subject of controversial debate. Before we offer an interpretation that situates the current set of findings within these literatures, we will briefly describe the predominant interpretations of the N2/P3 complex in both stopping and unexpected-event processing. In the realm of action-stopping, there is relatively widespread agreement on the fact that stopping involves a sequence of attentional detection of the stop-signal, followed by the implementation of motor inhibition (Matzke et al., 2013; Verbruggen et al., 2014). The earliest neuroscientific studies of the SST have proposed that the fronto-central P3 could index the inhibitory process of this cascade (de Jong et al., 1990, cf. Huster et al., 2013, for a review). Indeed, the P3 shows several features that reflect straightforward predictions regarding the inhibitory process that are directly derived from the race model of the stop-signal task. Both its peak and its onset occur earlier on successful compared to failed stop-trials (Kok et al., 2004; Wessel & Aron, 2015) and its timing indexes stop-signal reaction time across subjects (Wessel & Aron, 2015; Huster et al., 2019). In line with this, much subsequent work has focused on the proposal that the N2, which precedes the P3, could reflect a process that relates to the attentional processing of the stop-signal itself, or the detection of the associated conflict between the initiated response and the requirement to stop (Donkers & van Boxtel, 2004; Azizian et al., 2006; Enriquez-Geppert et al., 2010; Smith et al., 2010; Groom & Cragg, 2015). In the realm of unexpected-event processing, the exact nature of the mental processes reflected in the N2 and P3 events has been subject to intense debate as well, with the entirety of the literature too numerous to discuss (see Folstein & Van Petten, 2008, and Polich, 2007, for reviews). However, the dominant view of the fronto-central N2 (specifically, the N2b) is similar to that found in the stop-signal literature, in that it is commonly assumed to reflect the overt attentional processing of the event or the associated conflict (Näätänen & Gaillard, 1983; Folstein & Van Petten, 2008; Larson et al., 2014). The fronto-central P3 (also known as the P3a) after unexpected results has more heterogeneous interpretations, which range from working memory updating (Polich, 2007) to the evaluation of stimulus novelty (Friedman et al., 2001) to the mobilization for action following significant stimuli (Nieuwenhuis et al., 2011). While these two literatures are historically largely separate, the finding that both N2/P3 complexes originate from the same neural source suggest that they may indeed reflect the same cascade of processing in both situations (i.e., after stop-signals and unexpected events) – i.e., that processes that take place after stop-signals are also automatically engaged by unexpected events. This is backed up by findings from other imaging domains, such as the finding that unexpected events lead to the suppression of the motor system (Wessel & Aron, 2013; Dutra et al., 2018; Novembre et al., 2018; Novembre et al., 2019), and that they engage the subcortical circuitry that is ostensibly underlying inhibitory motor control via the basal ganglia (Bočková et al., 2011; Wessel et al., 2016; Fife et al., 2017). If it is indeed the case that the fronto-central N2/P3 complex reflects the same cascade of processes after both stop-signals and unexpected events, the specific relationships between the N2 and the P3 and the observed suppression of the SSVEP in the current study could provide a potential rejoinder to this literature. Specifically, the fact that the single-trial N2 was related to the interruption of the SSVEP to the *attended* stimulus lends support to the proposal that this potential reflects an attentional orienting to a salient, or, in this case, unexpected stimulus. This is in line with many conceptualizations from both the existing stop-signal and unexpected-event literature (see above), which largely converge in their interpretation of the N2. Additionally, the relationship between the trial-to-trial amplitude of the P3 and the observed interruption of the SSVEP to the unattended stimulus suggest that indeed, the P3 may reflect the activity of a ‘global’, non-selective inhibitory network that interrupts active *motoric and mental* representations when the situational demands call for it (such as after unexpected events). This is in line with our own recent theory about the activity of this network (Wessel & Aron, 2017), as well as with the proposal that the stop-signal P3 in particular reflects the implementation of the inhibitory part of the processing cascade during action-stopping (de Jong et al., 1990; Kok et al., 2004). Finally, it also could provide a hint towards a specific (inhibitory) mechanism by which unexpected events could aid the updating of current working memory contents (Polich, 2007), in line with recent studies of the activity of this inhibitory control mechanism in the suppression of mnemonic representations (Chiu & Egner, 2014, 2015; Wessel et al., 2016; Castiglione et al., 2019).

In summary, we have used a newly designed experimental paradigm to demonstrate that unexpected, task-irrelevant sounds lead to a suppression of the neural representation of both attended and unattended stimuli. Moreover, we used independent component analysis and single-trial analyses of EEG to show that these interruptions are related to specific separate aspects of the neural response to unexpected events within a neural system for inhibitory control. These findings provide a crucial potential expansion of the operating range of a well-characterized neural mechanism for cognitive control, and provide key insights into the cascade of neural and psychological processing that leads to distraction.

## Acknowledgements

The authors would like to thank Cathleen Moore, Eliot Hazeltine, Andrew Hollingworth, and Kai Hwang for helpful discussions of this project, and Nathan Chalkley, Brynne Dochterman, Kylie Dolan, Isabella Dutra, Alec Mather, and Daniel Thayer for help with data collection.

